# Surface-Induced cAMP Signaling Requires Multiple Features of the *Pseudomonas aeruginosa* Type IV Pili

**DOI:** 10.1101/2022.05.13.491915

**Authors:** S.L. Kuchma, G.A. O’Toole

## Abstract

*Pseudomonas aeruginosa* type IV pili (TFP) are important for twitching motility and biofilm formation. TFP have been implicated in surface sensing, a process whereby surface-engaged cells up-regulate synthesis of the second messenger cAMP to propagate a signaling cascade leading to biofilm initiation and repression of motility. Here we show that mutations in PilA impairing proteolytic processing of the prepilin into mature pilin as well as the disruption of essential TFP components, including the PilC platform protein and PilB assembly motor protein, fail to induce surface-dependent cAMP signaling. We show that TFP retraction by surface-engaged cells is required to induce signaling, and that the retractile motor PilT is both necessary and sufficient to power surface-specific induction of cAMP. The PilU retraction motor, in contrast, is unable to support full signaling in the absence of PilT. Finally, while we have confirmed that PilA and PilJ interact by bacterial two-hybrid analysis, our data do not support the current model that PilJ-PilA interaction drives cAMP signaling.

**Importance:** Surface sensing by *P. aeruginosa* requires TFP. TFP play a critical role in the induction of the second messenger cAMP upon surface contact; this second messenger is part of a larger cascade involved in the transition from a planktonic to biofilm lifestyle. Here we show that TFP must be deployed and actively retracted by the PilT motor for the full induction of cAMP signaling. Furthermore, the mechanism whereby TFP retraction triggers cAMP induction is not well understood, and our data argues against one of the current models in the field proposed to address this knowledge gap.

## Introduction

Bacteria are able to live as members of multicellular communities known as biofilms. As a prerequisite for this surface-associated sessile lifestyle, bacterial cells have the ability to recognize contact with a suitable surface to initiate colonization (1–3). The transition from a free-living to a surface-attached status occurs via a complex process that has been termed ‘surface sensing’. For *Pseudomonas aeruginosa*, a key feature of this process is up-regulation of the second messenger cAMP by cells upon surface engagement (1, 3). Work by several groups has implicated the type IV pilus (TFP), a dynamic, flexible filament assembled on the cell surface, as key to this surface-induced cAMP response, however the precise role of TFP in this process is not well understood (4, 5). For example, one model posits that interaction of the TFP with a surface triggers a signal transduction cascade through the PilJ/Chp sensory signaling circuit (4, 5). This signaling event has been proposed to require interaction between PilJ and the TFP pilin, PilA (5). PilJ is a chemosensory protein that upon receiving a stimulus is thought to activate the ChpA histidine kinase protein to induce autophosphorylation, based on analogy to the *E. coli* flagellar chemotaxis paradigm [(6); for reviews see (7, 8)]. ChpA then transduces this signal by phosphorylation of PilH and PilG, response regulator proteins (9) that signal to CyaB, the primary adenylate cyclase (AC), which in turn synthesizes cAMP from ATP (10). CyaA is a second AC that ostensibly plays a lesser role in this process (10).

The downstream effector responsive to changes in cAMP levels is the transcription factor Vfr, which binds to cAMP to regulate expression of numerous genes impacting a wide variety of functions, including Type II and Type III secretion systems involved in virulence as well as flagellar motility and quorum sensing (10, 11). Vfr also upregulates expression of genes required for TFP biogenesis, thereby propagating a positive feedback cycle producing more TFP to reinforce surface commitment and signaling (10, 12). Additionally, *P. aeruginosa* encodes a phosphodiesterase enzyme, CpdA, that degrades cAMP, allowing for negative modulation or a resetting of this signaling pathway (13). Notably, surface contact-induced signaling via Vfr/cAMP is consistent with previous work demonstrating that surface-engaged *P. aeruginosa* shows enhanced virulence (14).

TFP are motorized filaments comprised of the pilin subunit PilA. The powerhouse for the pilus is the inner membrane motor subcomplex comprised of the platform protein PilC and the ATP-hydrolyzing motor proteins that energize assembly (PilB) and disassembly (PilT/PilU) of the pilus filament [for reviews see (15, 16)]. It is thought that the PilC protein plays an integral role in transducing the chemical energy of ATP hydrolysis by the motors into the mechanical energy to drive both polymerization and depolymerization of pilin subunits in a coordinated manner (15, 16). Rapid cycling of polymerization, cell-surface attachment and depolymerization is responsible for a form of surface motility known as twitching, akin to a rope and grappling hook mechanism of movement (17). Pilus retraction also facilitates infection by bacteriophage that utilize the TFP as a receptor, with the retracting TFP bringing the phage into close proximity to the bacterial cell body (18).

Given the complexity of TFP assembly/disassembly and its intricate and over-lapping roles in surface sensing, twitching motility and surface attachment, we sought to explore different facets of TFP biology to assess their impact on surface sensing. Our findings here delineate the basic mechanistic requirements for the TFP in surface sensing by examining the pilin subunit as well as factors involved in assembly and function of the TFP. However, we did not find evidence to support a model in which the proposed PilA/PilJ interaction is the driving force for cAMP signaling, indicating that alternative mechanisms by which the TFP transmits a surface-engagement signal should be explored.

## Results

### PilA surface localization is important for surface-dependent induction of cAMP

In a previous study, we reported that soon after engaging a solid surface, *P. aeruginosa* responds by increasing production of the intracellular second messenger cAMP (4). This surface induction requires the PilA protein, the pilin subunit of the type IV pilus, as well as the PilJ/Chp signal transduction pathway (4). As a follow-up to those studies, we sought to more fully investigate the role of PilA and type IV pilus (TFP) biology in surface-dependent cAMP induction.

It has been well-established that the PilA protein undergoes two posttranslational modification events prior to pilus assembly. PilA is produced as a prepilin containing a short N-terminal leader peptide that is cleaved by the PilD peptidase between an invariant glycine (G^-1^) in the leader sequence and phenylalanine (F^+1^), the first residue of the mature protein (Fig. 1A), which is then methylated by PilD (19–21). Early genetic studies showed that cleavage of the leader peptide is a prerequisite for pilus assembly, whereas amino terminal methylation is not absolutely required and its precise role in pilus assembly remains unclear (19).

**Figure 1.**
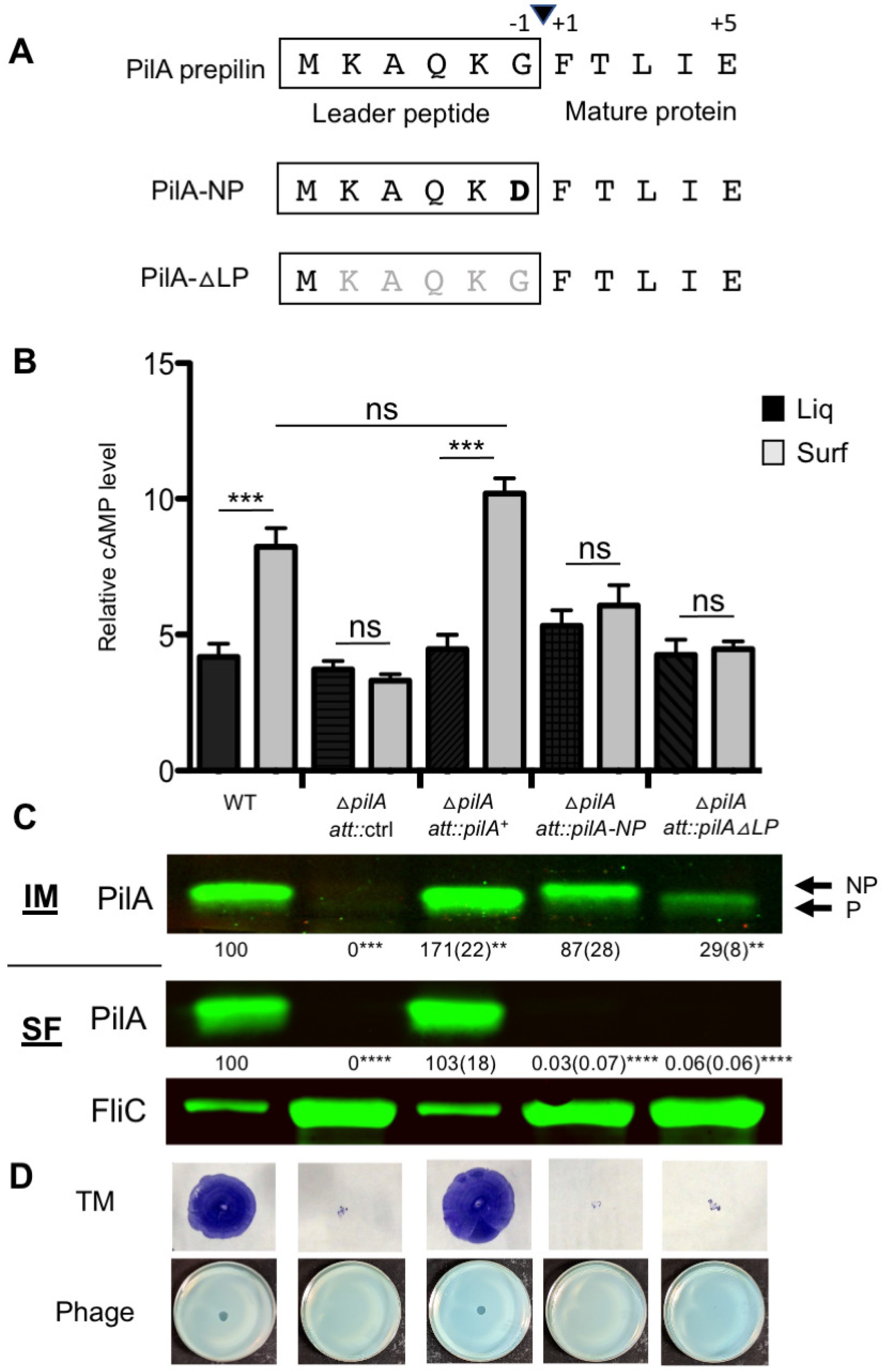
PilA surface localization is important for cAMP surface induction. **A)** Schematic showing the PilA leader peptide amino acid sequence (boxed) with cleavage site shown (black arrowhead) between the glycine in the −1 position (G^-1^) and the phenylalanine in the +1 position (F^+1^) of the mature protein. Shown below are 2 mutant variants: *pilA*-NP, a non-processed mutant with a substitution of the G^-1^ with an aspartic acid (D^-1^) and *pilA*-ΔLP, a deletion of the leader peptide and cleavage site (amino acids in gray). **B)** Relative cAMP levels in strains carrying the P_1_-*lacZ* cAMP reporter plasmid measured using β-gal assays as described in the Materials and Methods. The Δ*pilA* mutant strains contain an insertion of either the control sequence lacking a *pilA* allele or the indicated *pilA* allele at the chromosomal phage attachment (*att*) site (see Materials and Methods for details). Strains were grown in M8 minimal salts medium supplemented with glucose, MgSO4, CAA as well as 0.4 % arabinose (to induce expression of the *pilA* alleles) in either liquid (black bars) or on plates solidified with 1% agar (gray bars) and harvested for β-gal assays after 5 h of growth. Error bars indicate standard deviations of the average of three experiments with 2 replicates per strain and condition per experiment. Data were analyzed by one-way ANOVA followed by Tukey’s post-test comparison. ns, not significantly different; ***, *P* < 0.001. **C)** Western blot analysis of proteins isolated from strains grown on M8 1% agar plates as for the cAMP assay in panel B. The top panel shows PilA protein detected in the inner membrane (IM) fraction of cells, middle panel shows PilA levels in the sheared surface protein fraction (SF) and the bottom panel shows the FliC loading control. PilA protein was detected using an anti-PilA antibody and FliC with an anti-FliC antibody, and Western blots were imaged using the Odyssey CLx Imager and quantified using Image Studio Lite software. Levels of PilA in the inner membrane fraction (IM, top panel) were quantified relative to a loading control (non-specific cross-reacting band, not shown) and mean values (with standard deviations in parentheses) of three independent experiments shown below the IM panel with the WT set at 100%. Asterisks denote a significant difference of the indicated value compared to the WT; **, *P* < 0.01. PilA protein in the sheared surface protein fraction (SF, middle panel) is shown relative to the FliC protein (bottom panel) which serves as a loading control for this fraction. Samples in lanes 2, 4 and 5 were loaded at greater than five-fold higher levels (as determined by the relative FliC protein level) to confirm that no detectable PilA protein was present in these mutant fractions. Asterisks denote a significant difference of the indicated value compared to the WT; ****, *P* < 0.0001. **D)** Top panel shows images of twitching motility zones stained with crystal violet (CV) to aid in visualization. Twitching motility assays were performed using M8 1% agar plates containing 0.4 % arabinose and incubated for 24 h at 37°C, followed by 5 days at RT, at which point the agar was removed for CV staining. Bottom panel shows representative images of phage plaque assays performed by spotting a phage suspension onto an M8 plate seeded with a soft top agar (0.5%) layer inoculated with the indicated strain. Plates were incubated for 16 h at 37°C.

We first considered whether mutations that alter or prevent cleavage of PilA in the inner membrane, and thus prohibit proper surface localization of PilA, impact cAMP signaling. To test these ideas, we generated two mutations in the PiLA leader peptide that were previously shown to prevent cell surface pilus assembly (19). The first mutation is an amino acid substitution of the invariant G^-1^ residue with an aspartic acid (D) which prevents cleavage of the leader peptide by the PilD peptidase (Fig. 1A, referred to as *pilA*-NP, for not processed). The second mutation is a deletion of the leader peptide (LP) sequence excluding the initiator methionine and we refer to this mutant as *pilA*-ΔLP. These mutant *pilA* alleles as well as the wild-type (WT) *pilA^+^* allele were cloned into a mini-Tn7 element for single-copy insertion into the chromosomal *att* site of the Δ*pilA* deletion strain. These alleles are under control of the P_BAD_ arabinose-inducible promoter for expression at this site.

To assess the impact of these mutations on the cell’s ability to induce cAMP upon surface growth, we constructed a Δ*pilA* mutant strain carrying the P_1_-*lacZ* reporter construct and introduced the WT and various *pilA* alleles into the *att* site of this strain. The cAMP-responsive P_1_ promoter driving expression of the *lacZ* gene serves as a readout for cAMP levels, as previously described (4, 22). Thus, we could assess both cAMP signaling and the ability of the *pilA* mutant alleles to complement PilA function.

As shown in Figure 1B, we find that in contrast with the WT, the Δ*pilA att::ctrl* strain, which does not harbor a functional copy of *pilA* at the *att* site, fails to induce cAMP production upon surface growth relative to the wild-type strain (gray bars), consistent with previous work (4). Expression of the wild-type *pilA* allele at the *att* site restores cAMP surface induction, as expected for complementation of the Δ*pilA* mutant (Fig. 1B). We observe that neither strain expressing the leader peptide mutants from the *att* site in the Δ*pilA* mutant is capable of surface-induced cAMP production (Fig. 1B).

Given that the PilA protein resides in inner membrane pools prior to pilus assembly (19), we examined whether these PilA variant proteins are expressed and localized in the inner membrane fractions of surface-grown cells as compared to the WT and the complemented strain. As shown in Figure 1C (IM, top panel), the complemented Δ*pilA::att-pilA^+^* strain exhibits elevated levels of PilA protein in the inner membrane relative to the WT strain, as indicated by the values shown below each image, expressed as the percentage of WT protein levels (set at 100%) with standard deviation in parentheses. For the PilA-NP variant protein, we find that levels of this protein in the inner membrane are reduced by about 10% relative to the WT strain, although this difference is not statistically significant (*P* = 0.9) after correcting for multiple comparisons (Fig. 1C). Levels of the PilA-ΔLP protein variant are reduced approximately 70% relative to the WT PilA protein in the inner membrane, indicating that the PilA-ΔLP variant is perhaps less stable than the PilA-NP protein. As expected, the PilA-NP protein is larger in size than the mature processed PilA protein due to retention of the leader peptide, whereas the PilA-ΔLP protein is equivalent in size to the processed PilA proteins, given that this variant lacks the leader peptide by deletion rather than proteolytic cleavage.

To confirm that these mutations prevent PilA surface localization and pilus assembly, we examined the surface localized PilA protein, as a proxy for surface assembled pili, by isolating sheared surface protein fractions from these strains. As seen in the middle panel of Figure 1C (SF, surface fraction), we observe comparable levels of PilA protein in the Δ*pilA att::pilA^+^* complemented strain relative to the WT. Here, the flagellar filament FliC protein serves as a loading control for the surface protein fraction (bottom panel). For the Δ*pilA att::pilA-NP* and Δ*pilA att::pilA*Δ*LP* strains, we are unable to detect PilA on the surface of these cells, despite loading more than five-fold excess of the surface protein fractions (as evident by the elevated level of the FliC protein detected in these fractions), indicating that the PilA-NP and PilA-ΔLP proteins are unable to assemble into surface pili, consistent with previous findings (19).

To further verify these results, we performed twitching motility (TM) and phage sensitivity assays to assess the presence of functional cell surface pili for these strains. Twitching motility requires repeated rounds of coordinated pilus extension, attachment to the surface followed by retraction with force enough to pull the cell body towards the point of pilus attachment. As shown in Figure 1D, the top panel of images indicate that only the WT and the strain complemented with WT PilA are able to engage in twitching motility, and these strains show equivalent levels of twitching, confirming the complementation of the *pilA* mutation.

We also performed phage infection assays with phage DMS3vir (23), which allow for detection of less robust pilus function in comparison to TM, given that phage attachment and retraction to draw the phage to the cell body likely requires less force. The *pilA* leader peptide mutants are also deficient in any detectable pilus function because they are fully resistant to phage lysis, as is the *pilA* null mutant, compared to the WT and the complemented strain carrying the wild-type PilA (Fig. 1D, bottom panel). Together, these data indicate that the mutant PilA proteins tested here are not properly processed, not assembled into functional surface pili, and unable to initiate cAMP signaling in response to surface engagement.

### Mutations in conserved regions of PilA impact surface-induced cAMP signaling

To further explore the impact of PilA biology on the cAMP surface response, we generated additional mutations in the highly conserved alpha helical N-terminus of the PilA protein. We selected residues that have been implicated in important PilA functions, including pilus assembly and auto-regulation of *pilA* expression (19, 24–26). Figure 2A shows a schematic of the N-terminal amino acid sequence, including the leader peptide, of the PilA protein with the glutamate (E5) and proline (P22) residues highlighted in bold, and with the amino acid substitutions indicated in the boxes below. We quantified cAMP levels in the mutant strains and found that the only strain that fully retained the cAMP surface response was the Δ*pilA att::pilA-P22A* mutant (Fig. 2B, far right). In contrast, substitutions in the E5 residue, particularly those that alter the negative charge of this residue (E5A and E5K) show no surface response. Interestingly, the Δ*pilA att::pilA-E5D* mutant grown on a surface does show a modest increase in cAMP level relative to that of its liquid grown counterpart, although this difference is not statistically significant (*P* = 0.15) after correcting for multiple comparisons.

**Figure 2.**
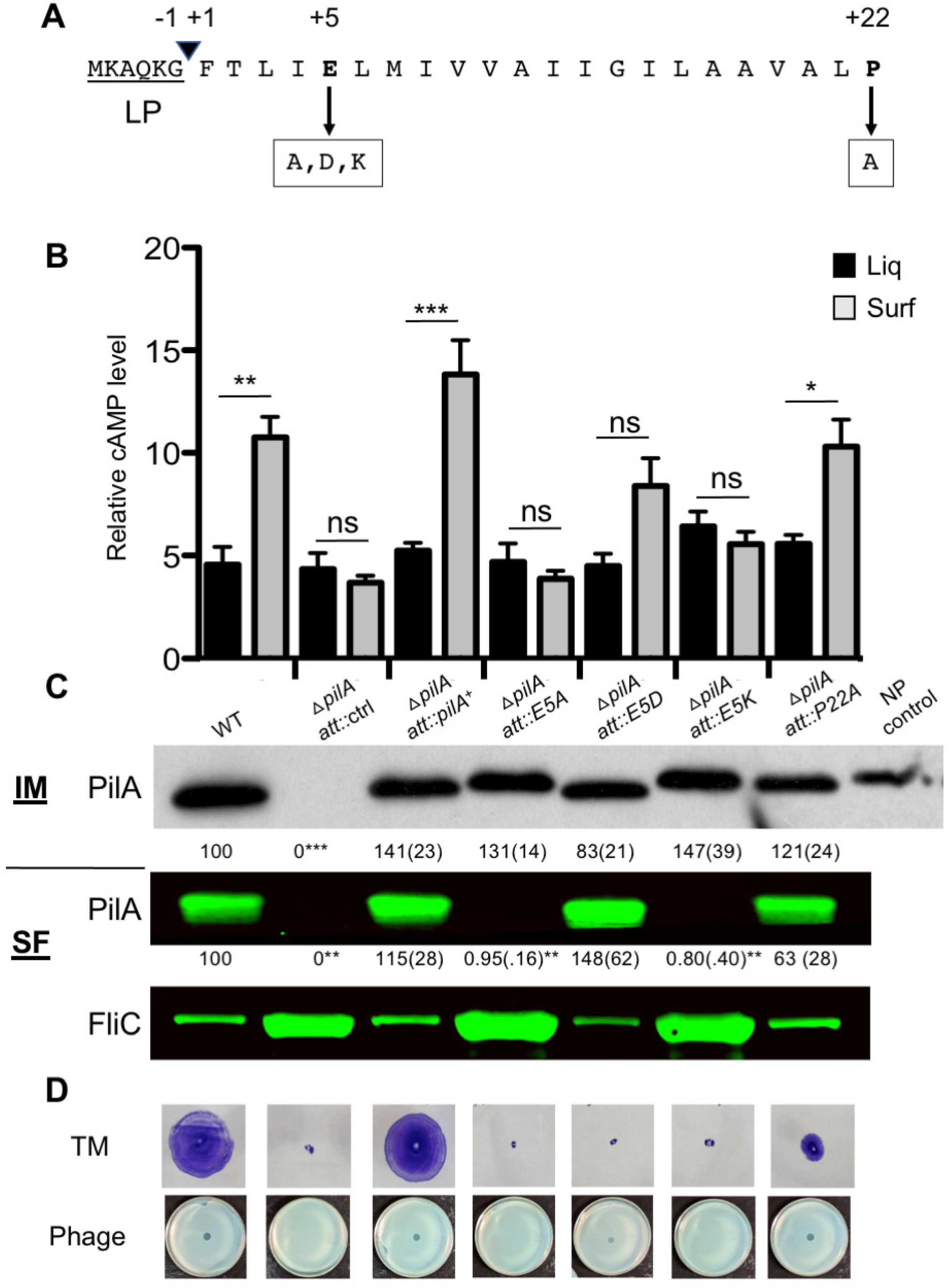
Surface pilus assembly is not sufficient for full cAMP signaling. **A)** Diagram of the N-terminal 22 amino acids of the PilA protein, including the leader peptide (underlined) with the arrowhead indicating cleavage position. The E5 and P22 residues (bold) were mutated to the indicated amino acids (boxed residues below the arrow) to determine the impact on cAMP surface signaling. **B)** The graph depicts cAMP quantification in strains carrying the P_1_-*lacZ* plasmid by β-gal assays. The Δ*pilA* mutant strains harbor an insertion of either the control sequence or the indicated *pilA* allele at the chromosomal phage attachment (*att*) site. Strains were grown in M8 medium with supplements as described (see Materials and Methods) and 0.4 % arabinose in either liquid (black bars) or on 1% agar plates (gray bars). Error bars indicate standard deviations of the average of three experiments with 2 replicates per strain and condition per experiment. Data were analyzed by one-way ANOVA followed by Tukey’s posttest comparison. ns, not significantly different; *, *P* < 0.05, **, *P* < 0.01, ***, *P* < 0.005. **C)** Western blot showing PilA in the inner membrane fractions (IM, top panel), the sheared surface fraction (SF, middle panel) and the FliC control (bottom panel). Western blots were developed with either the Clarity ECL kit (Bio-Rad; IM panel) or the Odyssey CLx Imager and quantified using Image Studio Lite software. PilA protein levels in the inner membrane fraction (IM, top panel) were quantified relative to a loading control (non-specific cross-reacting band, not shown) and mean values (with standard deviations in parentheses) of three independent experiments are shown below the IM panel with the WT set at 100%. Asterisks denote a significant difference of the indicated value compared to the WT; ***, *P* < 0.005. PilA protein in the sheared surface protein fraction (SF, middle panel) is shown relative to the the FliC protein as the loading control (bottom panel). Samples in lanes 2, 4 and 6 were loaded at least five-fold higher levels (as determined by the relative FliC protein level) to confirm that no detectable PilA protein was present in these mutant fractions. Asterisks denote a significant difference of the indicated value compared to the WT; **, *P* < 0.01. **D)** Top panel shows images of twitching motility zones stained with crystal violet (CV); bottom panel shows phage infection assays, both as described in Figure 1 legend.

When we assessed the levels of PilA protein in the IM fraction of surface-grown cells of these strains, we observed that all of the PilA protein variants are present at levels comparable to the WT and Δ*pilA att:: pilA^+^* complemented strain (Fig. 2C, top panel with protein quantifications below represented as % of WT), indicating the lack of surface cAMP induction is not due to the mutations impacting protein stability. Unexpectedly, we noted that the PilA-E5A and PilA-E5K variants are shifted in size relative to wild-type PilA protein and correspond to the size of unprocessed PilA-NP (IM panel, right-most lane), indicating that these variants likely did not undergo cleavage of the leader peptide. In previous genetic studies, the conserved E5 residue was shown to be important for N-methylation but did not impact leader peptide cleavage (19, 26). It is unclear why there is a discrepancy in these findings, but it is possible that strain differences may play some role as previous studies were performed with the PAO1 strain and we are using the *P. aeruginosa* PA14 strain here.

Consistent with our findings for unprocessed PilA from Figure 1, we observe here that the PilA-E5A and E5K variants are absent from surface protein fractions (Fig. 2C, second panel), despite loading greater than a five-fold excess of these fractions as in Figure 1. For the Δ*pilA att::pilA-E5A* and Δ*pilA att::pilA-E5K* mutants, we see both phage resistance and a loss of TM as expected for these strains that lack surface pili (Fig. 2D). In contrast, the PilA-E5D variant is detected at the processed size for mature PilA and is present in the surface protein fraction at levels comparable to the WT and complemented strain, indicating its assembly into surface pili (Fig. 2C, top two panels). Given that glutamate and aspartate are quite similar in terms of structure, size and charge, it is not surprising that this substitution is not fully disruptive to pilus assembly. Despite having surface pili, however, the Δ*pilA att::pilA-E5D* mutant is not capable of TM but is sensitive to phage infection, indicating that this mutant retains partial pilus functionality (Fig. 2D) in addition to residual cAMP signaling (Fig. 2B). The Δ*pilA att::pilA-P22A* mutant strain is unique among the mutants presented in Figures 1 and 2 in that it has surface pili that are functional in both TM, albeit to a level markedly below the WT (Fig. 2 and Supp. Fig. 1), and phage assays and shows a significant cAMP surface response.

Together, our observations here suggest that pilus assembly alone is not sufficient for robust cAMP signaling, given that the *pilA*-E5D mutant assembled surface PilA but did not show significant cAMP surface induction. That is, these data suggest pilus functionality (i.e., the ability to extend/retract) may play a key role in cAMP surface induction, a point we address below.

### PilC is required for surface-dependent induction of cAMP

To further address the notion that pilus assembly is required for cAMP signaling, we examined the cAMP surface response in a Δ*pilC* mutant, a strain lacking the inner membrane-anchored PilC protein shown to be essential for pilus biogenesis (27). PilC also serves as the structure onto which the ATPase motors assemble (27). As shown in Figure 3A, the Δ*pilC* mutant (Δ*pilC att::*ctrl) is defective for cAMP surface induction compared to the WT and Δ*pilA att:: pilA^+^* complemented strains.

**Figure 3.**
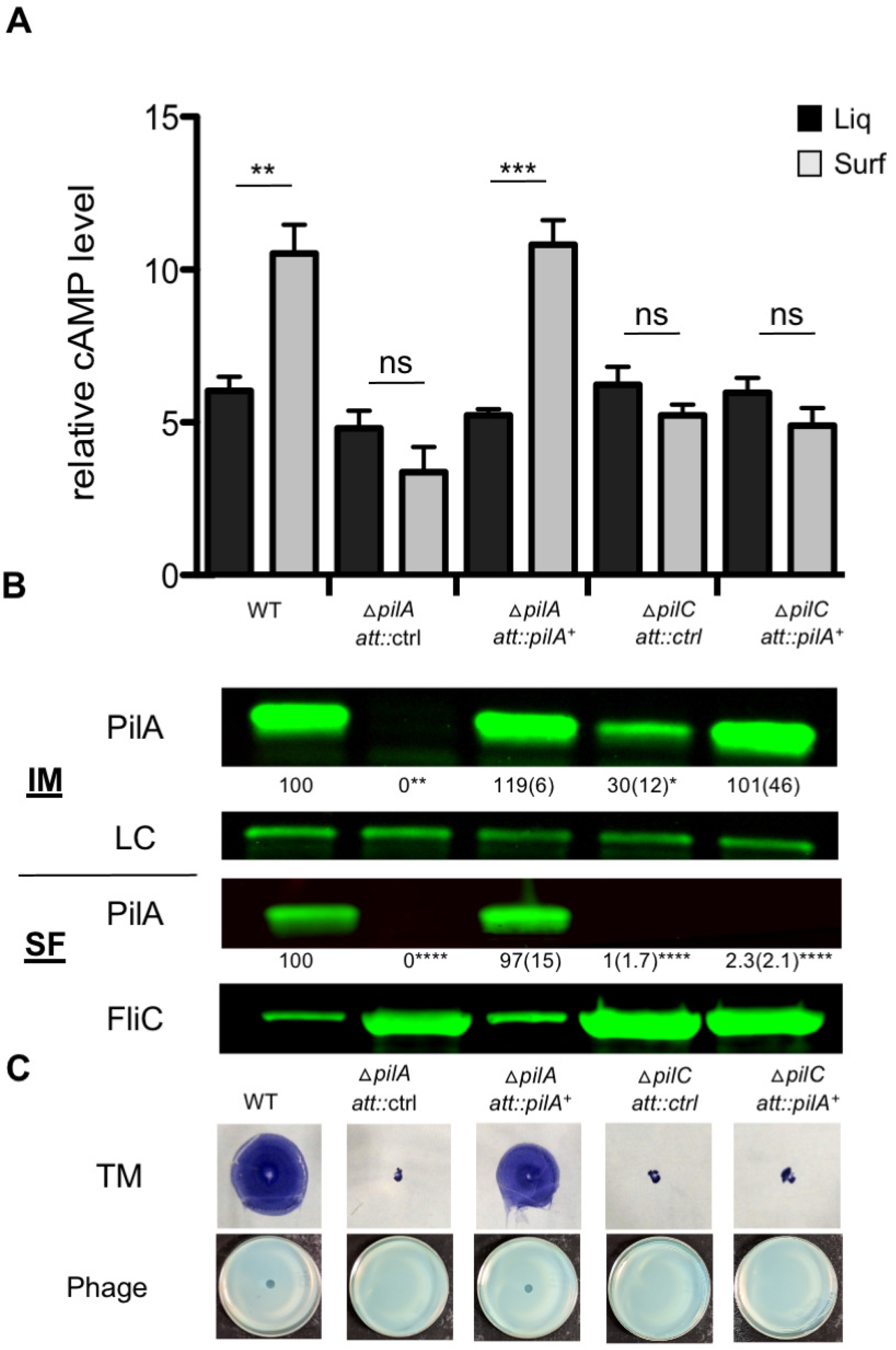
Loss of the PilC platform protein prevents pilus assembly and cAMP signaling. **A)** The graph shows relative cAMP levels in strains carrying the P_1_-*lacZ* reporter plasmid measured using β-gal assays as described in the Material and Methods. Strains were grown in M8 medium with supplements as described (see Material and Methods) and 0.4 % arabinose in either liquid (black bars) or on 1% agar plates (gray bars). Error bars indicate standard deviations of the average of three experiments with 2 replicates per strain and condition per experiment. Data were analyzed by one-way ANOVA followed by Tukey’s post-test comparison. ns, not significantly different; **, *P* < 0.01, ***, *P* < 0.001. **B)** Western blot showing PilA protein in cellular fractions of surface-grown cells. Levels of PilA in the inner membrane fraction (IM, top panel) were quantified relative to the loading control (lower panel, LC) and the mean values (with standard deviations in parentheses) of three independent experiments are shown with the WT at 100%. Asterisks denote a significant difference relative to the WT; *, *P* < 0.05 **, *P* < 0.01. PilA protein in the sheared surface protein fraction (SF, 3^rd^ panel) is shown relative to the FliC protein (4^th^ panel) as the loading control for this fraction. Samples in lanes 2, 4 and 5 were loaded at more than five-fold higher levels to confirm that no detectable PilA protein was present in these mutant fractions. Asterisks indicate a significant difference relative to the WT; ****, *P* < 0.0001. Proteins were detected using polyclonal antibodies to PilA and FliC and Western blots were imaged using the Odyssey CLx Imager and quantified using Image Studio Lite software. **C)** Top panel shows images of twitching motility zones stained with crystal violet (CV). Bottom panel shows phage infection assays.

We also examined PilA protein levels in this mutant by Western blotting of cellular fractions prepared from surface-grown cells. In the inner membrane (IM) fraction which serves as a cellular reservoir for the PilA monomer, we observed an ~70% decrease in PilA protein levels in the Δ*pilC att::*ctrl strain relative to the WT and complemented Δ*pilA* strains (Figure 3B, top panel). In the surface protein fraction (SF), we did not observe the PilA protein in the *pilC* mutant, a result that was expected given the critical role of the PilC protein in pilus assembly.

To rule out that the lack of cAMP stimulation was due to a general reduction in levels of PilA in the *pilC* mutant, we inserted a copy of the *pilA* gene *in trans* at the *att* site of the Δ*pilC* mutant, and assessed the resulting Δ*pilC att::pilA^+^* strain by Western blotting. The results showed PilA protein levels in the Δ*pilC att::pilA^+^* strain were restored to levels comparable to the WT strain in the IM fraction (Figure 3B, values below top panel), but despite this increase, we did not observe PilA protein in the SF fraction (Figure 3B, 3^rd^ panel) nor any cAMP surface signaling (Figure 3A) for the Δ*pilC att::pilA^+^* strain. As shown in Figure 3C, we further confirmed that the Δ*pilC* mutant is unable to engage in TM (top row) and is resistant to phage infection (bottom row) regardless of whether PilA levels are enhanced or not. These data further support the conclusion that, at a minimum, pilus assembly is a requirement for proper surface cAMP induction.

### Surface pili are necessary but not sufficient for cAMP surface induction

Our data above suggest that TFP assembly may be necessary, but not sufficient for surface-induced cAMP signaling. To further address the question of whether and to what degree pilus function is required for cAMP signaling, we turned to the motor proteins responsible for powering pilus movement. Pilus extension and retraction are the critical features that help drive twitching motility, phage sensitivity and other important aspects of pilus biology. Pilus extension occurs via pilus assembly and is powered by an ATPase motor protein, PilB, which catalyzes the polymerization of PilA monomers into pilus filaments using energy from ATP hydrolysis. Pilus retraction occurs by disassembly or depolymerization of pili into pilin monomers and retraction is powered primarily by the action of the ATPase motor protein PilT. A third PilT-like ATPase motor protein, PilU, also displays features consistent with a role in retraction and indeed, several recent studies have shown that PilU is an ATPase, but that its role in retraction is dependent upon the presence of PilT (28, 29). Moreover, the dependence of PilU function on the presence of PilT was shown in different bacterial species, including *P. aeruginosa*, and may represent a paradigm for other organisms that possess both retractile motor homologs (28, 29). The impacts of these findings for *P. aeruginosa* on our experiments will be further addressed below when we consider the role of the retraction ATPases in cAMP surface signaling.

We first addressed the role of the PilB assembly motor in the cAMP surface response. The graph in Figure 4A shows that the Δ*pilB* mutant is not capable of inducing cAMP production in response to surface growth. As expected, this mutant strain is defective for TM (Fig. 4B, top row) and is resistant to phage infection (Fig. 4B, 2^nd^ row), indicative of a lack of surface pili. We confirmed this lack of pili by Western blotting of PilA in surface protein fractions (Fig. 4C, SF). However, this strain is capable of synthesizing the PilA protein and importantly, we do observe the pilin monomer in the IM, the reservoir for pilus assembly, in this strain (Fig. 4C, IM).

**Figure 4.**
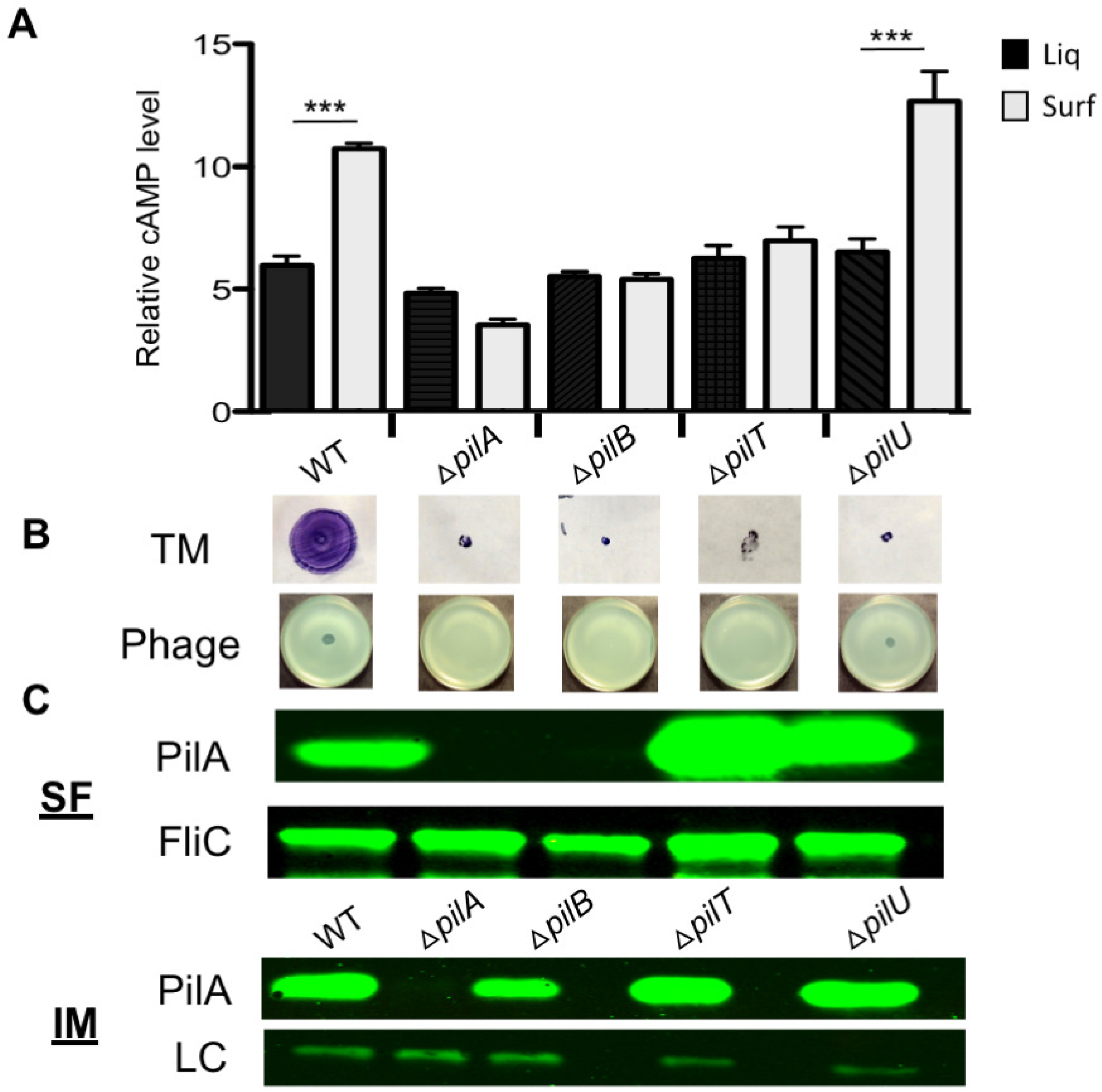
Pilus motor function is required for cAMP signaling. **A)** The graph shows relative cAMP levels in strains carrying the P_1_-*lacZ* plasmid measured using β-gal assays as described in the Materials and Methods. Strains were grown in M8 medium in either liquid (black bars) or on plates solidified with 1% agar (gray bars). Error bars indicate standard deviations of the average of three experiments with 2 replicates per strain and condition per experiment. Data were analyzed by one-way ANOVA followed by Tukey’s post-test comparison. ***, *P* < 0.001. **B)** Phenotypic assays of motor mutants. Top panel shows images of CV-stained twitching motility zones. Twitching motility assays were performed by inoculating M8 1% agar plates and incubating for 24 h at 37°C, followed by 5 days at RT, and CV staining. Bottom panel shows images of phage plaque assays, performed by spotting a phage suspension onto an M8 plate seeded with a soft top agar layer inoculated with the indicated strain. Plates were incubated for 16 h at 37°C. **C)** Top two panels show the sheared surface fraction (SF) from the indicated strains. PilA protein was detected using an anti-PilA antibody. The FliC protein serves as a loading control in this fraction (lower panel) and was detected using an anti-FliC antibody. Bottom panels, IM fraction isolated from whole cell lysates of the strains shown. A non-specific band detected in all samples serves as a loading control (LC). Western blots were developed using IR-Dye^®^-labeled fluorescent secondary antibodies and imaged using the Odyssey CLx Imager.

In considering the retractile motors, PilT and PilU, here we built upon data that has recently been reported by our group and others (4, 5, 30) to synthesize a more thorough picture of the role these motors play in surface induction of cAMP. As was reported by Lee et al. (30), we also observed that Δ*pilT* mutants are unable to respond to surface growth by inducing cAMP production (Fig. 4A). In contrast, the Δ*pilU* mutant remains fully capable of initiating cAMP signaling in response to surface growth and shows an elevated level of cAMP surface induction relative to the wild-type strain, indicating the possibility that PilU may have a negative impact on cAMP surface stimulation. In terms of surface pilus assembly and function, the Δ*pilT* mutant, which is capable of pilus assembly but not retraction, exhibits elevated levels of surface pili (Fig. 4C), consistent with previous findings (31). The over-abundance of surface pili is attributed to the defect in pilus retraction as the Δ*pilT* mutant produces wild-type levels of the PilA monomer in IM fractions of this mutant strain (Fig. 4C, IM panel). Despite the excess of surface pili, the Δ*pilT* mutant is defective for TM and resistant to phage infection (Fig. 4B, C) (31), highlighting the importance of PilT-driven retraction in these phenotypes.

The Δ*pilU* mutant is also hyper-piliated, as seen by elevated PilA protein levels in the surface fraction of these strains (Fig. 4C), and defective for TM, but in contrast to the Δ*pilT* mutant, the Δ*pilU* mutant remains sensitive to phage infection (Fig. 4B) (9, 31, 32). What these results indicate is that the Δ*pilU* mutant retains some measure of retraction ability, likely due to the presence of a functional PilT in this strain. Given that the most notable differences between the Δ*pilT* and Δ*pilU* mutants are the cAMP surface response and phage sensitivity, together these findings suggest that surface pili are necessary but not sufficient for cAMP surface stimulation. Rather, some level of pilus retraction when cells are on a surface is required for this phenomenon to occur (note that the *pilU* mutant does not show cAMP signaling when grown in liquid), which is in general agreement with our findings regarding the *att::pilA-E5D* and *att::pilA-P22A* mutants described above.

As mentioned above, several recent studies have illuminated the dependence of PilU function in pilus retraction on the presence of PilT. The implication of these data is that, in a Δ*pilT* mutant, the PilU motor is unable to contribute to retraction, effectively rendering the Δ*pilT* mutant a Δ*pilT ΔpilU* double mutant. Interestingly, the two studies in *V. cholerae* regarding the roles of PilT and PilU in retraction of the *Vibrio cholerae* Type IV DNA uptake pilus (28, 29) and MSHA pilus (28) both showed that PilU ATPase activity was essential for pilus retraction in either a *pilT*(K136A) Walker A (ATP-binding motif) or *pilT*(E204A) Walker B (ATP hydrolysis motif) ATPase non-functional mutant strain, indicating that PilU was able to power retraction as long as PilT was present, even if PilT lacked ATPase activity (28, 29).

Given these data, we wondered whether restoration of PilU function in a Δ*pilT* mutant would be able to rescue retraction ability and impact cAMP surface induction, indicating whether or not retraction by either ATPase was able to influence the cAMP surface response. To test this possibility, we expressed either a *pilT*^WA^ or a *pilT^WB^* ATPase mutant allele as compared to a wild-type *pilT* control allele in the Δ*pilT* mutant background and assessed the impact on TM, phage infection and cAMP surface induction. If PilU is able to power retraction in the presence of a non-functional PilT counterpart, we would expect to see some restoration of pilus function such as phage sensitivity and possibly TM. Indeed, as shown in Fig. 5A, we observe that the Δ*pilT att::pilT*^WA^ and Δ*pilT att::pilT^WB^* strains regain phage sensitivity similar to the complemented Δ*pilT att::pilT^+^* strain, and in contrast to the phage-resistant Δ*pilT att::control* parental strain.

**Figure 5.**
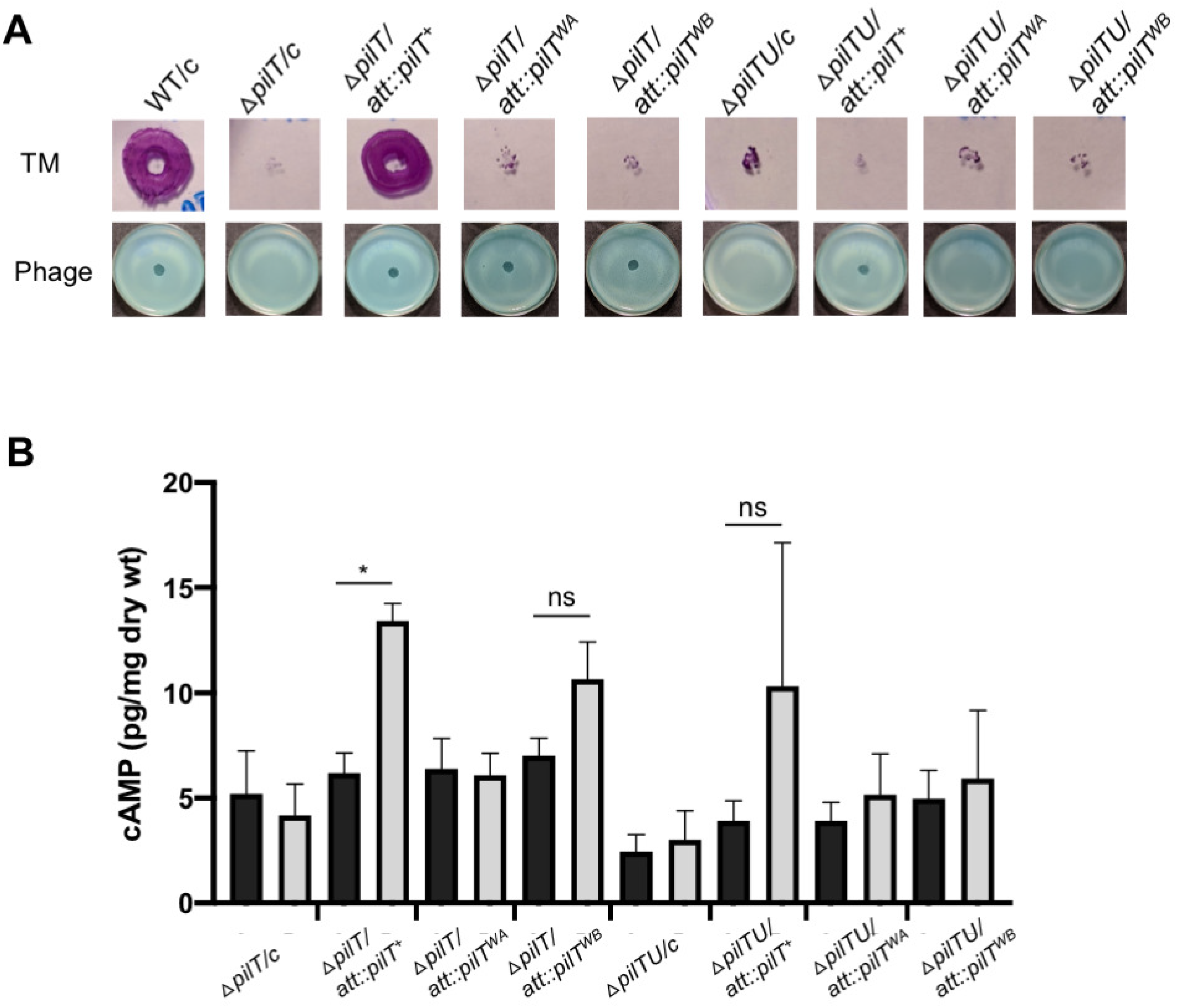
Phenotypes of Walker Box mutations in PilT. **A)** TM (top) and phage sensitivity (bottom) phenotypes of the indicated PilT alleles in the WT and the Δ*pilU* or Δ*pilTU* mutant backgrounds. Strains were grown and assayed as described in Materials and Methods, with 0.25% arabinose added for induction of *pilT* gene expression in each experiment. **B)** cAMP quantification in Δ*pilT* and Δ*pilTU* mutant strains expressing the indicated *pilT* alleles from the *att* site. The *pilT*^WA^ allele signifies the K136A mutation in the Walker A motif (involved in ATP binding) and the *pilT*^WB^ allele signifies the E204A mutation in the Walker B motif (involved in ATP hydrolysis).

To confirm that these *pilT*^WA^ and *pilT^WB^* alleles encode non-functional versions of PilT and are not simply complementing loss of PilT function, we expressed these alleles in a Δ*pilT* Δ*pilU* double mutant. The resulting Δ*pilT* Δ*pilU att:: pilT*^WA^ and Δ*pilT* Δ*pilU att:: pilT^WB^* strains exhibit resistance to phage infection, indicating that the phage sensitivity observed in the Δ*pilT att::pilT*^WA^ and Δ*pilT att::pilT^WB^* strains depends on restoring PilU function and not by complementing PilT function. However, restoration of PilU function in these strains does not rescue the twitching motility defect, indicating that PilU activity alone is not able to power pilus retraction sufficient for this behavior.

Given that our findings indicate PilU is active in the Δ*pilT att:: pilT*^WA^ and Δ*pilT att::pilT^WB^* strains, we next assessed whether PilU-mediated pilus retraction is able to trigger cAMP surface induction in the absence of a fully functional PilT protein (Figure 5B). These results show that the Δ*pilT att::pilT*^WA^ strain does not exhibit a significant cAMP surface response as compared to the Δ*pilT att::pilT^+^* strain, which shows the expected surface-induced increase in cAMP, indicating that the *pilT* Walker A mutant does not support PilU function in cAMP signaling. In the case of the Δ*pilT att::pilT^WB^* mutant, this strain exhibits a modest increase in cAMP level in surface-grown cells but this effect is not statistically significant after correcting for multiple comparisons (*P* = 0.825). Together, these data suggest that retraction by PilU is not sufficient to fully support a cAMP surface response in the absence of a functional PilT retraction motor, although PilU might allow some residual cAMP signaling in the absence of the WT PilT motor.

### Genetic studies do not support the model that PilJ-PilA interactions drive cAMP signaling

Our data are consistent with a model wherein PilA must be assembled into pili on the cell surface and those pili must retain some retraction activity to effectively initiate the cAMP cascade upon surface engagement. However, questions remain regarding the precise mechanisms by which pili engage the cAMP surface signaling response. Previous studies have shown interaction of PilA with the MCP PilJ via the bacterial two-hybrid assay (B2H), leading to a proposed model whereby PilA signals directly to PilJ to influence cAMP production via the Pil/Chp signaling cascade (5). The *pilJ* gene encodes a methyl-accepting chemotaxis protein (MCP) involved in signal transduction. When stimulated, the PilJ protein activates the ChpA kinase which in turn activates the response regulators, PilH and PilG, leading to cAMP production via the CyaB adenylate cyclase (33).

In light of this model, we examined several *pilA* mutants with varied impacts on cAMP levels and evaluated whether there was any alteration in interaction with PilJ using the B2H system. For example, the *pilA*(E5A) and *pilA*(E5K) mutants do not exhibit surface induction of cAMP relative to the wild type (Fig. 2); however, these PilA variants showed increased interaction with PilJ relative to the wild-type protein (Fig. 6A-C). Additionally, the PilA(P22A) variant interacts with PilJ better than the wild-type PilA protein and is produced at levels equivalent to the WT PilA in the B2H assay strain, but we do not observe an enhancement in cAMP surface induction in the *pilA*(P22A) strain (Fig. 6A-C, see also Fig. 2). Finally, the E5D mutation shows an equivalent level of interaction with PilJ compared to the WT, but shows reduced cAMP signaling (Fig. 6A-C, see also Fig. 2).

**Figure 6.**
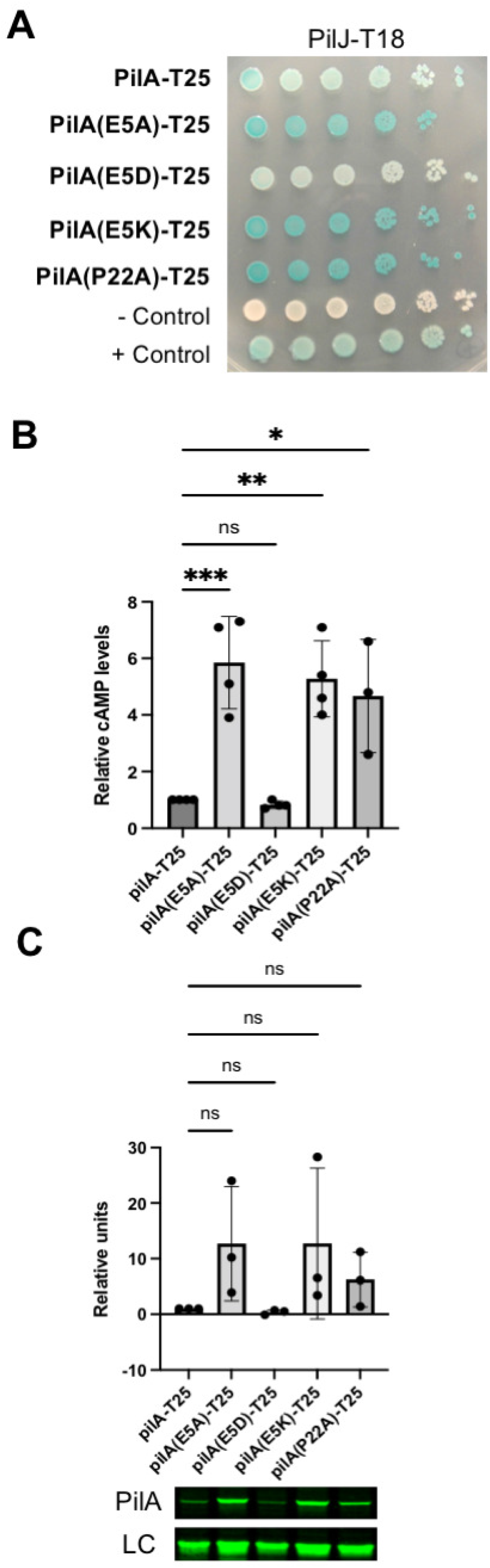
The extent of PilJ-PilA interaction does not correlate with observed cAMP signaling. **A)** Representative image of a bacterial two-hybrid assay plate indicating the interaction between the PilJ-T18 fusion protein with the PilA-T25 proteins listed on the left side of the plate. Plates were grown for 24h at 30°C. **B)** Graph showing quantification of B2H interactions. Bacterial cells were harvested from plates grown as described in **A** and β-galactosidase assays were performed as described in the Methods section. Error bars indicate standard deviations of the average of three experiments with 2 replicates per strain. Mean values for each interaction were normalized to the wild type PilA-T25/PilJ-T18 interaction in each experiment. Data were analyzed by one-way ANOVA followed by Tukey’s post-test comparison. ns, not significantly different; *, P < 0.05; **, P < 0.01; ***, *P* < 0.001. **C)** Graph showing quantification of PilA-T25 fusion protein levels in the B2H interactions from panel B. The panel below shows a representative image of the PilA-T25 fusion proteins detected for the indicated interaction using the anti-PilA antibody. A non-specific band detected in all samples serves as a loading control (LC, bottom panel). Western blots were developed using IR-Dye^®^-labeled fluorescent secondary antibodies, imaged using the Odyssey CLx Imager and quantified using Image Studio Lite software. Protein levels for each interaction were normalized to the wild type PilA-T25/PilJ-T18 interaction in each experiment. Error bars indicate standard deviations of the average of three experiments.

This lack of correlation between the B2H assay results and cAMP measurements could possibly be explained, at least in part, by the differences between the assay systems, but not by changes in stability of the mutant proteins (Fig. 6C). Furthermore, we have used changes in interaction between proteins in the B2H to effectively probe functional consequences of protein-protein interactions in previous studies (34–36). Thus, these discrepancies suggest that a PilA-PilJ interaction may not be functionally important for surface-dependent cAMP induction and indicate there might be additional or alternative mechanisms at play which remain to be discovered.

## Discussion

Here we examined features of TFP biology to better understand the role of this appendage in surface sensing. We show that mutations blocking cleavage of the prepilin or deletion of the PilA leader sequence eliminates surface-induced cAMP signaling. We interpret these data to mean that assembly of pili on the cell surface is required for detecting surface engagement. An alternative interpretation is that the process of cleavage or engagement of pilin with the leader peptidase enzyme (PilD) is specifically required for signaling to occur. However, analysis of the *pilC* platform mutant and the *pilB* assembly motor mutant, which are both capable of making mature, processed PilA protein but are unable to support pilus assembly and cannot engage in surface signaling, argue against this interpretation. Instead, the phenotypes of these assembly-specific mutants provide further support for the conclusion that surface assembly of pili is necessary for cAMP signaling.

By examining a second class of pili mutants – those that can assemble surface pili – for pilus function and surface signaling, we concluded that pilus assembly is necessary but not sufficient for signaling. Rather, some level of pilus function (i.e., extension and retraction) is required. The clearest example of this distinction came from comparing the retraction motor mutants, Δ*pilT* and Δ*pilU*. The Δ*pilT* mutant is hyper-piliated but unable to engage in TM and is resistant to phage infection, indicating lack of retraction. In contrast, the Δ*pilU* mutant, while also hyper-piliated, and TM negative, is sensitive to phage infection and thereby active for retraction. Only the Δ*pilU* mutant is capable of surface-specific cAMP signaling. These data are consistent with a model whereby the minimal requirement for pilus function in signaling is assembly plus retraction that is sufficient to support phage infection but not twitching motility. The presumed reduced level of force required for phage sensitivity versus twitching motility calls into question whether the forces required to power motility are needed for cAMP signaling, a finding which can inform model building.

Our results led to the question of whether both retractile motors PilT and PilU are capable of generating force to support surface sensing. A simple interpretation of the differences between the Δ*pilT* and Δ*pilU* mutants might lead one to conclude that only the PilT motor is capable of powering surface signaling, given that the Δ*pilU* mutant retains a functional PilT motor and only the Δ*pilU* mutant is competent for retraction and signaling. However, based on recent data showing the dependence of PilU function on the presence of PilT, and thus reasoning that the Δ*pilT* mutant was likely to be a Δ*pilT* Δ*pilU* double mutant, we tested whether PilU was able to support retraction and signaling as the sole retraction motor in the presence of PilT mutants with non-functional ATPase activity. Our data indicate that while PilU does regain the ability to support phage infection and therefore retraction under these conditions, this motor does not exhibit robust surface cAMP induction. Thus, it may be that PilT is uniquely suited to engage in surface signaling. Furthermore, in light of observations that loss of PilU correlates with an elevated baseline level of surface-specific cAMP stimulation, it seems reasonable to propose that PilU might serve to modulate the ability of PilT to engage in surface signaling, possibly as a means to fine-tune the cAMP surface response. Consistent with this idea, a recent study from our team examining cell-surface engagement of *P. aeruginosa* over time shows that prior surface exposure primes cells for enhanced surface engagement upon subsequent surface exposure in a cAMP-dependent manner, and furthermore, Δ*pilU* mutant cells show a marked acceleration of this “surface adaptation” process relative to wild-type cells (30). However, we cannot rule out the possibility that this accelerated response may, in part, be due to the excess surface pili produced by the *pilU* mutant. Studies aimed at understanding the functions and dynamics of these retraction motors in surface signaling are ongoing.

Our data are consistent with models proposed by others suggesting that tension on the pilus is involved in signaling surface engagement (5, 37). This tension is likely due to surface engagement of the pilus coupled with retraction of surface-engaged pili. However, the question of how retraction of a surface-engaged pilus fiber transduces this signal to the cell’s interior to stimulate a surface response, and the amount of force required to trigger such signaling, remains a challenging one to answer. In *P. aeruginosa*, an observed interaction between PilA and PilJ has led to the prediction that this PilA-PilJ interaction plays a role in driving surface sensing. However, we did not find evidence to support this model. Instead, we observed a lack of correlation between the impact of PilA variants on cAMP surface induction and the ability of these variants to interact with PilJ.

How then might pilus retraction trigger signaling? While our studies here do not support PilA-PilJ interaction as the driving force for signaling, the notion that PilA may directly interact with a signaling partner is compelling given that such an interaction between PilA and a signaling partner to influence surface sensing was recently highlighted in studies in *Caulobacter crecentus* where interaction of PilA with the inner membrane-localized sensor kinase PleC is proposed to couple surface attachment of this bacterium with cell cycle progression, an important aspect of the *C. crescentus* lifecycle (38). The PilA segment required for interaction with PleC has been defined by bacterial two-hybrid analysis to be a 17 amino acid peptide within the alpha helical trans-membrane domain that includes the highly conserved E5 residue of PilA and is referred to as CIP (cell cycle initiating peptide) for its role in regulating cell cycle progression (38). To explain how the PilA CIP mediates surface sensing in *C. crescentus*, the authors propose that surface attachment of pili and retraction by depolymerization of pilus filaments leads to accumulation of PilA monomers in the inner membrane and this accumulation leads to increased interaction of CIP with a transmembrane region of PleC. Interaction of PleC with PilA is then proposed to activate PleC’s kinase activity in phosphorylation of downstream factors involved in promoting cell cycle progression. A similar mechanism for sensing PilA levels in the inner membrane has been proposed in *P. aeruginosa* to explain the observed inverse relationship between inner membrane levels of PilA subunits and *pilA* gene transcription, whereby high levels of PilA inner membrane reservoirs correlate with a decrease in *pilA* gene expression and vice versa (25). This regulatory feedback mechanism to monitor and adjust levels of pilin produced by cells is reported to involve direct interaction of the PilA protein with the sensor kinase PilS that, together with its cognate response regulator PilR, decreases *pilA* gene expression (25). Notably, this study and the *C. crescentus* CIP work may represent an emerging theme in pilus biology – that is, the direct participation of pilin monomers in signal transduction.

In the study pertaining to PilA feedback regulation on PilSR in *P. aeruginosa*, two *pilA* mutations in particular, E5K and P22A, disrupted both the interaction of PilA with PilS as well as *pilA* gene expression. These two amino acids are notable for their high level of conservation across Type IV pilins and their roles in pilus assembly. The E5 residue of the mature pilin protein imparts a negative charge in the N-terminal alpha helix of the pilin monomer that is proposed to engage in electrostatic interaction with the positive charge of the F1 residue of the adjacent pilin subunit in the growing pilus filament to facilitate polymer assembly (24, 39). The P22 residue is believed to introduce a kink in the alpha helix, aiding in destabilizing a segment of the helix and allowing this alpha helix to unfold and pack into the pilus core during pilus assembly (24, 39). As we described above, the E5K charge reversal mutation eliminates cAMP surface signaling, however, this result is due to a defect in pre-pilin cleavage which precludes its role in pilus assembly. For a P22A mutation, in our study, this mutant makes surface pili that support phage infection and also TM but with a greatly reduced zone of TM relative to that for the WT, suggesting that a proline in this position is not essential for pilus assembly/retraction but is important for optimal function. Furthermore, we observe that the P22A mutant allows for surface signaling, indicating that this proline residue, and perhaps by extension the helix kink, is not necessary for the cAMP surface response. Moreover, this result suggests that the pilin sensing mechanism reported by Kilmury et. al. (25), and the surface sensing mechanism examined here likely represent distinct signaling phenomena. Further studies are needed to fully address this possibility.

In this report our findings highlight key features of the PilA pilin and TFP function required for surface sensing by *P. aeruginosa*. We identified mutations that render the cells defective for TM, but still capable of retaining some pilus function as judged by phage sensitivity. These mutations still retain surface-dependent cAMP signaling. Together, these findings suggest that TM per se is *not* essential for surface sensing, an important observation as we continue to build models to explain TFP-dependent surface sensing. Finally, the finding that a functional PilT is required for PilU-dependent cAMP signaling on the surface indicates that surface sensing requires not only pilus retraction but a specific role for PilT in this mechanism. Overall, this work has further enhanced our understanding of factors required to, as well as dispensable for, surface-dependent cAMP signaling.

## Acknowledgements

The work was supported by funding from NIH Grant R37 AI83256 to G.A.O. We also received support from the Bio-MT Molecular Tools and Molecular Interactions and Imaging Core (P20-GM113132).

## Materials and Methods

### Strains and Media

Strains used in this study are listed in Supplemental Table S1. *P. aeruginosa* PA14, *E. coli* DH5α, S17-1 λpir, BTH101, JM109, and AR31110 were routinely cultured in lysogeny broth (LB) medium, solidified with 1.5% agar when necessary. Gentamicin (Gm) was used at 25 μg/ml for *P. aeruginosa* and at 10 μg/ml for *E. coli*. Carbenicllin (Cb) was used at 250 μg/ml for PA and 100 μg/ml for *E. coli*. Nalidixic acid (NA) at 20 μg/ml, streptomycin at 100 μg/ml, and kanamycin (Kan) at 50 μg/ml for *E. coli*. For phenotypic assays with *P. aeruginosa*, M8 minimal salts medium (as indicated) were supplemented with MgSO_4_ (1mM), glucose (0.2%), and casamino acids (CAA; 0.5%). For expression plasmids harboring the *P_BAD_* promoter, arabinose was added to cultures at either 0.2% or 0.4% final concentration, as noted. To visualize ß-galactosidase activity derived from co-expression of the bacterial two-hybrid constructs, 5-bromo-4-chloro-3-indolyl-β-D-galactopyranoside (X-Gal; 40 μg/ml) and isopropyl-D-thiogalactopyranoside (IPTG; 0.5 mM) was added to selective plates. *S. cerevisiae* strain InvSc1 (Invitrogen), used for plasmid construction via *in vivo* homologous recombination, was grown with yeast extract-peptone-dextrose (1% Bacto yeast extract, 2% Bacto peptone, and 2% dextrose), as reported. Selections with InvSc1 were performed using synthetic defined agar-uracil (Qbiogene, 4813-065).

### Construction of mutant strains and plasmids

Supplemental Table S2 lists all plasmids used in this study. Primers used in plasmid construction and mutant construction are listed in Supplemental Table S3. In-frame gene deletions were performed via allelic exchange, as previously described (40). Plasmids for this purpose were constructed via cloning by homologous recombination of relevant PCR products into the pMQ30 vector in yeast, as reported (41) or using Gibson assembly. For integration of single copy genes on the *P. aeruginosa* PA14 chromosome at the *att* site, original mini-Tn7 vectors (42) or a modified version (pMQ56-mTn7) allowing for cloning by homologous recombination in yeast were utilized. Integration of mini-Tn7 elements was performed as reported (42). When necessary, the Gm-resistance cassette was removed from the mini-Tn7 via the flanking FRT sites using the pFLP2 plasmid followed by *sacB-*mediated plasmid counterselection as reported (43). Point mutations and small deletions in the *pilA* gene were generated using either primer-based mutations in PCR or QuikChange^®^ site-directed mutagenesis followed by homologous recombination of mutated PCR products in yeast or by Gibson assembly. Constructs for plasmid-based expression of genes were generated using PCR and Gibson Assembly^®^ (NEB, Boston, MA) followed by cloning into pMQ72. For all plasmids and constructs used in the experiments described herein, the relevant cloned genes were fully sequenced to confirm that the correct sequences were present.

### Motility assays

Twitching motility plates were prepared using M8 medium supplemented with glucose, MgSO_4_ and CAA and 0.4% arabinose, where indicated, and solidified with 1% agar. TM plates were inoculated as previously described, followed by incubation for 24 h at 37°C and for 5 days at room temperature thereafter. To visualize the twitch zones, the agar was removed and stained by addition of 0.1 % crystal violet (CV) to the plate. Images were taken and twitch zones measured using ImageJ. Swarm motility plates were prepared with M8 medium supplemented with glucose, MgSO_4_ and CAA and 0.4% arabinose and solidified with 0.5 % agar. Swarm assays were performed as previously described (44). Swim motility plates were prepared with M8 medium supplemented with glucose, MgSO_4_ and CAA and 0.4% arabinose and solidified with 0.3 % agar. Swim assays were performed as previously described (45).

### Phage plaque assays

For phage plaque assays, small (60 x 15 mm) plates were prepared using M8 medium supplemented with glucose, MgSO_4_ and CAA and solidified with 1% agar; arabinose (0.4 %) was added where indicated. To seed the bacterial lawn, a 1 ml solution of molten 0.5 % agar (M8 media supplemented as for the plates) was prepared for each plate and inoculated with the appropriate *P. aeruginosa* strain (50 ul from fresh overnight LB liquid culture), mixed and then poured on the plate, making sure to cover entire plate surface. To inoculate with phage, a 2 ul aliquot of a DMS3 lytic phage suspension (DMS3_vir_;(23)) was spotted on the solidified top agar layer of the plate and allowed to dry. The plate was then incubated for 16 h at 37°C.

### β-galactosidase assays for cAMP quantification

For strains harboring either the pR-P_1_-*lacZ* cAMP reporter fusion or pR-*lacZ* control plasmid (33), overnight LB-grown cultures were diluted 1:100 in regular M8 liquid medium and grown to OD_600_ of ~ 0.4 (mid-log phase) at 37 °C with vigorous shaking. At this point, an aliquot from the liquid culture (200 ul) was spread to M8 1% agar plates and both liquid cultures and plates were incubated at 37°C for 5 hours. Cells were then harvested from liquid cultures or scraped from the agar surface. β-galactosidase reactions were performed as previously described (4).

### Bacterial two-hybrid (B2H) assays

Protein-protein interactions were evaluated using the published <u>b</u>acterial <u>a</u>denylate <u>c</u>yclase two-<u>h</u>ybrid (BATCH) system obtained from Euromedex (Souffelweyersheim, France) (46, 47). In this study, the wild-type *pilA* gene or *pilA* mutant alleles were cloned into the pKT25 vector and tested in interactions using the pilJ-*T18* construct generated previously (4) by co-transformation into *E. coli* BTH101 cells. For visualization of the interactions, transformants were 10-fold serially diluted and spotted (2 ul) on LB agar containing Cb, Kan, X-Gal (5-bromo-4-chloro-3-indolyl-β-D-galactopyranoside) (40 g/ml), and IPTG (isopropyl-D-thiogalactopyranoside) (0.5 mM) and incubated for at least 24 h at 30°C. To quantify interaction efficiency, transformations were diluted and plated to LB agar plates containing Cb, Kan and IPTG and cells were harvested after 24 h growth at 30°C and subjected to *β*-galactosidase activity assays as previously described (4).

### Statistical analysis

The Prism software package (GraphPad, San Diego, CA) was used for statistical analysis of experimental data.

### Protein detection and cellular localization experiments

Bacterial strains were grown either in M8 liquid cultures or on M8 1% agar plates for harvest and isolation of proteins for Western blotting, corresponding to the conditions of growth used for cAMP quantification via β-gal assays described above. Whole cell lysates (WC) and cellular fractions were prepared as previously described (48). Total protein concentrations in whole cell and inner membrane fractions were quantified using the Pierce™ BCA protein assay kit (ThermoFisher Scientific, Waltham, MA). Sheared surfaced fractions were prepared as previously described (49) with the modification that BSA (1 mg/ml final concentration in PBS) was added during precipitation with PEG and NaCl to facilitate protein recovery. For Western blotting, equivalent total protein quantities from WCL and IM samples and equal volumes of sheared surface fractions were resolved by SDS-PAGE using either 18% (Figure 1) or 12% polyacrylamide gels (all other experiments). Proteins transferred to a nitrocellulose membrane were probed with either anti-PilA or anti-FliC antisera. The FliC protein is the subunit of the *Pseudomonas* polar flagellum and serves as a loading control for the sheared surface protein fractions. Detection of proteins via Western blotting was performed using either the Clarity ECL detection kit (Bio-Rad, Hercules, CA) or by fluorescence detection using IR-Dye^®^-labeled fluorescent secondary antibodies and imaged using the Odyssey CLx Imager (LICOR Biosciences, Inc., Lincoln, NE), as indicated. Protein quantifications were performed using Image Studio Lite software (LICOR Biosciences, Inc., Lincoln, NE). Relative protein levels were calculated by subtracting the signal in the Δ*pilC att::*ctrl sample lane as background, normalized to a loading control and expressed as a percentage of the WT signal set to 100%.

## References

1. G. A. O’Toole, G. C. Wong, Sensational biofilms: surface sensing in bacteria. Curr. Opin. Microbiol. 30, 139–146 (2016).

2. R. Belas, Biofilms, flagella, and mechanosensing of surfaces by bacteria. Trends Microbiol. 22, 517–527 (2014).

3. B.-J. Laventie, U. Jenal, Surface sensing and adaptation in bacteria. Annu. Rev. Microbiol. 74, 735–760 (2020).

4. Y. Luo, et al., A hierarchical cascade of second messengers regulates *Pseudomonas aeruginosa* surface behaviors. MBio 6, e02456–14 (2015).

5. A. Persat, Y. F. Inclan, J. N. Engel, H. A. Stone, Z. Gitai, Type IV pili mechanochemically regulate virulence factors in *Pseudomonas aeruginosa*. Proc. Natl. Acad. Sci. U. S. A. 112, 7563–7568 (2015).

6. C. B. Whitchurch, et al., Characterization of a complex chemosensory signal transduction system which controls twitching motility in *Pseudomonas aeruginosa*. Mol. Microbiol. 52, 873–893 (2004).

7. G. H. Wadhams, J. P. Armitage, Making sense of it all: bacterial chemotaxis. Nat. Rev. Mol. Cell Biol. 5, 1024–1037 (2004).

8. M. A. Matilla, D. Martín-Mora, J. A. Gavira, T. Krell, *Pseudomonas aeruginosa* as a model to study chemosensory pathway signaling. Microbiol. Mol. Biol. Rev. 85, e00151–20 (2021).

9. J. J. Bertrand, J. T. West, J. N. Engel, Genetic analysis of the regulation of type IV pilus function by the Chp chemosensory system of Pseudomonas aeruginosa. J. Bacteriol. 182, 194–1010 (2010).

10. M. C. Wolfgang, V. T. Lee, M. E. Gilmore, S. Lory, Coordinate regulation of bacterial virulence genes by a novel adenylate cyclase-dependent signaling pathway. Dev. Cell 4, 253–263 (2003).

11. F. E.L., et al., The *Pseudomonas aeruginosa* Vfr regulator controls global virulence factor expression through cyclic AMP-dependent and -independent mechanisms. J. Bacteriol. 192, 3553–3564 (2010).

12. S. A. Beatson, C. B. Whitchurch, J. L. Sargent, R. C. Levesque, J. S. Mattick, Differential regulation of twitching motility and elastase production by Vfr in *Pseudomonas aeruginosa*. J. Bacteriol. 184, 3605–3613 (2002).

13. E. L. Fuchs, et al., In vitro and in vivo characterization of the *Pseudomonas aeruginosa* cyclic AMP (cAMP) phosphodiesterase CpdA, required for cAMP homeostasis and virulence factor regulation. J. Bacteriol. 192, 2779–2790 (2010).

14. A. Siryaporn, S. L. Kuchma, G. A. O’Toole, Z. Gitai, F. M. Ausubel, Surface attachment induces *Pseudomonas aeruginosa* virulence. Proc. Natl. Acad. Sci. U. S. A. 111, 16860–16865 (2014).

15. L. L. Burrows, *Pseudomonas aeruginosa* twitching motility: Type IV pili in action. Annu. Rev. Microbiol. 66, 493–520 (2012).

16. L. Craig, K. T. Forest, B. Maier, Type IV pili: dynamics, biophysics and functional consequences. Nat Rev Micro 17, 429–440 (2019).

17. D. E. Bradley, A function of *Pseudomonas aeruginosa* PAO polar pili: twitching motility. Can. J. Microbiol. 26, 146–154 (1980).

18. D. E. Bradley, Evidence for the retraction of *Pseudomonas aeruginosa* RNA phage pili. Biochem. Biophys. Res. Commun. 47, 142–149 (1972).

19. M. S. Strom, S. Lory, Amino acid substitutions in pilin of *Pseudomonas aeruginosa.*Effect on leader peptide cleavage, amino-terminal methylation, and pilus assembly. J. Biol. Chem. 266, 1656–1664 (1991).

20. D. N. Nunn, S. Lory, Product of the *Pseudomonas aeruginosa* gene pilD is a prepilin leader peptidase. Proc. Natl. Acad. Sci. U. S. A. 88, 3281–3285 (1991).

21. J. C. Pepe, S. Lory, Amino acid substitutions in PilD, a bifunctional enzyme of Pseudomonas aeruginosa. Effect on leader peptidase and N-methyltransferase activities in vitro and in vivo. J. Biol. Chem. 273, 19120–19129 (1998).

22. N. B. Fulcher, P. M. Holliday, E. Klem, M. J. Cann, M. C. Wolfgang, The *Pseudomonas aeruginosa* Chp chemosensory system regulates intracellular cAMP levels by modulating adenylate cyclase activity. Mol. Microbiol. 76, 889–904 (2010).

23. K. C. Cady, J. Bondy-Denomy, G. E. Heussler, A. R. Davidson, G. A. O’Toole, The CRISPR/Cas adaptive immune system of *Pseudomonas aeruginosa* mediates resistance to naturally occurring and engineered phages. J. Bacteriol. 194, 5728–5738 (2012).

24. F. Wang, et al., Cryoelectron microscopy reconstructions of the pseudomonas aeruginosa and *Neisseria gonorrhoeae* Type IV Pili at sub-nanometer resolution. Structure 25, 1423–1435 (2017).

25. S. L. N. Kilmury, L. L. Burrows, Type IV pilins regulate their own expression via direct intramembrane interactions with the sensor kinase PilS. Proc. Natl. Acad. Sci. U. S. A. 113, 6017–22 (2016).

26. B. L. Pasloske, W. Paranchych, The expression of mutant pilins in *Pseudomonas aeruginosa:* fifth position glutamate affects pilin methylation. Mol. Microbiol. 2, 489–495 (1988).

27. H. K. Takhar, K. Kemp, M. Kim, P. L. Howell, L. L. Burrows, The platform protein is essential for type IV pilus biogenesis. J. Biol. Chem. 288, 9721–9728 (2013).

28. D. W. Adams, J. M. Pereira, C. Stoudmann, S. Stutzmann, M. Blokesch, The type IV pilus protein PilU functions as a PilT-dependent retraction ATPase. PLoS Genet. 15, e1008393 (2019).

29. J. L. Chlebek, et al., PilT and PilU are homohexameric ATPases that coordinate to retract type IVa pili. PLoS Genet. 15, e1008448 (2019).

30. C. K. Lee, et al., Multigenerational memory and adaptive adhesion in early bacterial biofilm communities. Proc. Natl. Acad. Sci. U. S. A. 115, 4471–4476 (2018).

31. C. B. Whitchurch, M. Hobbs, S. P. Livingston, V. Krishnapillai, J. S. Mattick, Characterisation of a *Pseudomonas aeruginosa* twitching motility gene and evidence for a specialised protein export system widespread in eubacteria. Gene 101, 33–44 (1991).

32. C. B. Whitchurch, J. S. Mattick, Characterization of a gene, pilU, required for twitching motility but not phage sensitivity in *Pseudomonas aeruginosa*. Mol. Microbiol. 13, 1079–1091 (1994).

33. N. B. Fulcher, P. M. Holliday, E. Klem, M. J. Cann, M. C. Wolfgang, The *Pseudomonas aeruginosa* Chp chemosensory system regulates intracellular cAMP levels by modulating adenylate cyclase activity. Mol. Microbiol. 76, 889–904 (2010).

34. A. E. Baker, et al., Flagellar stators stimulate c-di-GMP production by *Pseudomonas aeruginosa*. J. Bacteriol. 201, e00741–18 (2019).

35. K. M. Dahlstrom, K. M. Giglio, A. J. Collins, H. Sondermann, G. A. O’Toole, Contribution of physical interactions to signaling specificity between a diguanylate cyclase and its effector. MBio 6, e01978–15 (2015).

36. S. S. Webster, C. K. Lee, W. C. Schmidt, G. C. L. Wong, G. A. O’Toole, Interaction between the type 4 pili machinery and a diguanylate cyclase fine-tune c-di-GMP levels during early biofilm formation. Proc. Natl. Acad. Sci. U. S. A. 118, e2105566118 (2021).

37. C. K. Ellison, et al., Obstruction of pilus retraction stimulates bacterial surface sensing. Science 358, 535–538 (2017).

38. L. Del Medico, D. Cerletti, P. Schächle, M. Christen, B. Christen, The type IV pilin PilA couples surface attachment and cell-cycle initiation in *Caulobacter crescentus*. Proc. Natl. Acad. Sci. 117, 9545–9553 (2020).

39. L. Craig, et al., Type IV pilin structure and assembly: X-ray and EM analyses of Vibrio cholerae toxin-coregulated pilus and *Pseudomonas aeruginosa* PAK pilin. Mol. Cell 11, 1139–1150 (2003).

40. H. P. Schweizer, Allelic exchange in *Pseudomonas aeruginosa* using novel ColE1-type vectors and a family of cassettes containing a portable *oriT* and the counter-selectable *Bacillus subtilis sacB* marker. Mol. Microbiol. 6, 1195–1204 (1992).

41. R. M. Q. Shanks, N. C. Caiazza, S. M. Hinsa, C. M. Toutain, G. A. O’Toole, Saccharomyces cerevisiae-based molecular tool kit for manipulation of genes from gram-negative bacteria. Appl. Environ. Microbiol. 72, 5027–5036 (2006).

42. K. H. Choi, H. P. Schweizer, mini-Tn7 insertion in bacteria with single attTn*7* sites: Example *Pseudomonas aeruginosa*. Nat. Protoc. 1, 153–161 (2006).

43. T. T. Hoang, R. R. Karkhoff-Schweizer, A. J. Kutchma, H. P. Schweizer, A broad-host-range Flp-FRT recombination system for site-specific excision of chromosomally-located DNA sequences: application for isolation of unmarked *Pseudomonas aeruginosa* mutants. Gene 212, 77–86 (1998).

44. D.-G. Ha, S. L. Kuchma, G. A. O’Toole, Plate-based assay for swarming motility in *Pseudomonas aeruginosa*. Methods Mol Bio 1149, 67–92 (2014).

45. D.-G. Ha, S. L. Kuchma, G. A. O’Toole, Plate-Based assay for swimming motility in *Pseudomonas aeruginosa*. Methods Mol Bio 1149, 59–65 (2014).

46. G. Karimova, J. Pidoux, A. Ullmann, D. Ladant, A bacterial two-hybrid system based on a reconstituted signal transduction pathway. Proc. Natl. Acad. Sci. U. S. A. 95, 5752–5756 (1998).

47. A. Battesti, E. Bouveret, The bacterial two-hybrid system based on adenylate cyclase reconstitution in *Escherichia coli*. Methods 58, 325–334 (2012).

48. S. L. Kuchma, E. F. Griffin, G. A. O’Toole, Minor pilins of the type IV pilus system participate in the negative regulation of swarming motility. J. Bacteriol. 194, 5388–5403 (2012).

49. C. L. Giltner, M. Habash, L. L. Burrows, *Pseudomonas aeruginosa* minor pilins are incorporated into type IV Pili. J. Mol. Biol. 398, 444–461 (2010).

